# A novel proviral mechanism for the ISG, Shiftless, during human coronavirus infection

**DOI:** 10.64898/2026.10.16.759718

**Authors:** Taylor M. Shue, Georgios Mourtzouchos, Dar-Yin Li, Yee T. Ong, Antonis Athanasiadis, Maria Bousali, Madeleine K. Sanders, Sophie N. Spector, Christopher Rivera-Bonilla, Khadija Ndoye, Kenneth C. Levenson, Jens Cuba, Jayden Harris, Kelly Mieck, Shuiyun Lan, Volker Kinast, Julia LeCher, Jérémie Le Pen, Stefan G. Sarafianos, Eleftherios Michailidis

## Abstract

As highly pathogenic human coronaviruses continue to emerge from zoonotic sources, understanding how host intrinsic immune responses control infection is critical for preventing severe disease. The interferon (IFN) pathway induces antiviral effectors, including the interferon-stimulated gene (ISG) Shiftless (SHFL), which restricts −1 programmed ribosomal frameshifting during SARS-CoV-2 infection. Here, we investigate a novel proviral mechanism for SHFL using loss-of-function and gain-of-function approaches. Despite its established antiviral activity, SHFL enhanced infection by SARS-CoV-2 and the seasonal coronavirus HCoV-229E by increasing cellular susceptibility to infection. SHFL further promoted coronavirus infection by reducing cellular sensitivity to IFNα, through IFN signaling. Finally, we demonstrate that SHFL’s proviral effect is not completely dependent on the −1 programmed ribosomal frameshifting domain. In contrast, SHFL-mediated regulation of IFN signaling is independent of this domain. Together, these findings establish SHFL as a dual-function ISG with distinct proviral and antiviral activities, highlighting the complex roles of innate immune effectors during coronavirus infection.

**Importance:** Highly pathogenic human coronaviruses have emerged three times since 2002, and spillovers are likely to continue as human contact with animal habitats increases. Against a novel pathogen, the innate immune response is the host’s first line of defense, and interferon-stimulated genes (ISGs) are its main effectors. ISGs are canonically antiviral, but growing evidence suggests that viruses may co-opt them for proviral functions. Here we describe a novel, proviral function of the ISG, Shiftless (SHFL), during infection by SARS-CoV-2 and the seasonal coronavirus HCoV-229E. Our findings show that an IFN-induced gene is not necessarily an antiviral factor. For human coronaviruses, SHFL expression initially benefits the virus although it may act as an antiviral factor at a later step. Knowing which ISGs have dual functions, and in which cells, is a prerequisite for interpreting context-dependent IFN responses and for understanding the balance between host defense and viral persistence.

## Introduction

SARS-CoV, MERS-CoV, and SARS-CoV-2 each crossed into humans from animal reservoirs since 2002 and caused substantial human disease^1–4^. This rate of emergence of highly pathogenic coronaviruses is attributed to environmental changes and humans expanding into habitats where animal reservoirs of coronaviruses are present^5^. The most recent human coronavirus, SARS-CoV-2, caused the COVID-19 pandemic, which has resulted in over seven million deaths worldwide since its emergence in 2019^6^. Beyond mortality, SARS-CoV-2 is also associated with long-term consequences of infection (e.g., long COVID) and unprecedented impacts on society, culture, and the global economy^7^. Vaccination against SARS-CoV-2 has reduced severe disease and death, but it relies on adaptive immunity and its protection against infection wanes^8^. By contrast, the innate immune system requires no prior exposure to a pathogen to restrict the viral lifecycle. Understanding the virus-host interactions that govern this response and whether this response is broadly applicable to a diverse range of coronaviruses will inform future research into preventative and therapeutic measures against viruses that emerge in the future.

A central arm of the innate immune system is the type I interferon (IFN) response pathway^9^. During viral infection, pattern recognition receptors (PRRs) detect viral nucleic acids and activate signaling cascades that result in type I IFN production. Type I IFNs signal in an autocrine and paracrine manner to turn on the production of interferon-stimulated genes (ISGs)^10^. This establishes an antiviral state that helps control the magnitude and spread of infection^10,11^. For SARS-CoV-2, the RNA sensors retinoic acid-inducible gene I (RIG-I) and melanoma differentiation-associated gene 5 (MDA5) are the dominant molecules driving the IFN response^12^. Mutations that disrupt the proper production of, or response to, IFN are linked to the development of severe COVID-19 disease^13–15^. There are hundreds of ISGs, at least 14 of which restrict different stages of the SARS-CoV-2 lifecycle, including entry, replication, and assembly/egress^16,17^. This response is so potent that SARS-CoV-2 has evolved to antagonize the IFN pathway through multiple mechanisms^18^. Although ISGs are canonically considered antiviral effectors, there is a growing body of evidence showing that viruses can co-opt ISGs to help complete their lifecycle and that individual ISGs can have dual functions depending on their cellular context^16,19,20^. One example is the ISG lymphocyte antigen 6 family member E (LY6E)^21,22^. LY6E inhibits SARS-CoV-2 entry by interfering with membrane fusion, but it also aids influenza A virus by promoting uncoating after the viral particle is released from the endosome. Other factors that influence the role of an ISG include cell type and host species. Whether an ISG can be co-opted in this way during coronavirus infection, however, remains largely untested: most ISGs characterized for SARS-CoV-2 to date have been studied as restriction factors, with only a handful showing proviral activity during coronavirus infection^23–25^.

In our previously published unbiased genome-wide arrayed CRISPR knockout (KO) screen for SARS-CoV-2 host factors^26^, SHFL was detected as a potential host-dependency factor. This screen was performed in Huh-7.5 cells, a human hepatocellular carcinoma cell line. While the liver is not the primary tissue target of SARS-CoV-2, Huh-7.5 cells are naturally susceptible to the virus and have been widely used in SARS-CoV-2 research^27–33^. In addition to virus susceptibility, these cells are advantageous for the study of ISGs because they are incapable of robust IFN signaling through RIG-I, but responsive to exogenous IFN treatments to trigger JAK/STAT signaling and downstream ISG production^34^. This allows for precise regulation of IFN in the system and ability to probe ISG functionality during infection without IFN stimulation. SHFL is canonically an antiviral ISG with activity across a wide range of RNA and DNA viruses. SHFL restricts viral infection through a variety of mechanisms, largely centered on RNA binding to inhibit viral translation and reduce the stability of viral RNAs (vRNAs). Relevant to this study, SHFL has been shown to restrict human immunodeficiency virus type 1 (HIV-1) and SARS-CoV-2 by decreasing the efficiency of −1 programmed ribosomal frameshifting (−1 PRF), an essential step in translation of the viral replicase polyprotein^35–42^. In an independent study of SHFL’s antiviral role during flavivirus infection, Hanners et al. tested a panel of viruses in SHFL-overexpressing Huh-7.5 cells^43^. They showed SHFL overexpression increased infection with SARS-CoV-2 and the seasonal coronavirus HCoV-OC43, both of which are in the betacoronavirus genus; however, the underlying mechanism remained elusive. With this additional evidence, consistent with our screening data from an orthogonal approach, we hypothesized that SHFL may belong to the growing list of ISGs with both antiviral and proviral activity.

Here, we test this hypothesis using complementary loss-of-function and gain-of-function approaches. We generated clonal SHFL knockouts in Huh-7.5 cells and asked whether SHFL’s host-dependency phenotype holds for SARS-CoV-2 and the seasonal alphacoronavirus HCoV-229E, extending the question across two genera of the *Coronaviridae*. Next, we tested whether SHFL acts on the establishment of infection, the release of progeny virions, or both. We then overexpressed wild-type SHFL and a naturally occurring isoform lacking the −1 PRF domain in A549-ACE2-TMPRSS2 (A549-AT) cells, to ask whether the phenotype is specific to Huh-7.5 cells and whether it maps to the domain that carries SHFL’s antiviral activity. Finally, we combined a mixed-population infection assay with an IFNα dose-response and JAK/STAT inhibition to ask how the proviral activity of SHFL relates to the interferon response that induces it. Together, these findings establish SHFL as a dual-function ISG with distinct proviral and antiviral activities, highlighting the complex roles of innate immune effectors during coronavirus infection.

## Results

### Identification of the interferon-stimulated gene Shiftless as a candidate host dependency factor for SARS-CoV-2

Using a genome-wide arrayed CRISPR KO screen in Huh-7.5 cells to identify novel host factors for SARS-CoV-2, SHFL KO unexpectedly decreased the percentage of SARS-CoV-2 infected cells, indicating a potential proviral function for SHFL. This was based on immunofluorescence (IF) for SARS-CoV-2 nucleocapsid (N) protein (Fig. 1A)^26^. The infection data was normalized to account for variation in cell count and plate-to-plate differences and reported as a z-score as previously described^26^. The SARS-CoV-2 z-score for SHFL KO did not change in untreated cells but decreased under IFNα pre-treatment conditions. Our findings suggest that the phenotype may be IFN-dependent, like IFNAR1, but in the opposite direction. Under IFNα pre-treatment, SHFL functions as a host dependency factor similar to ACE2, the SARS-CoV-2 entry receptor (Fig. 1B).

**Figure 1.**
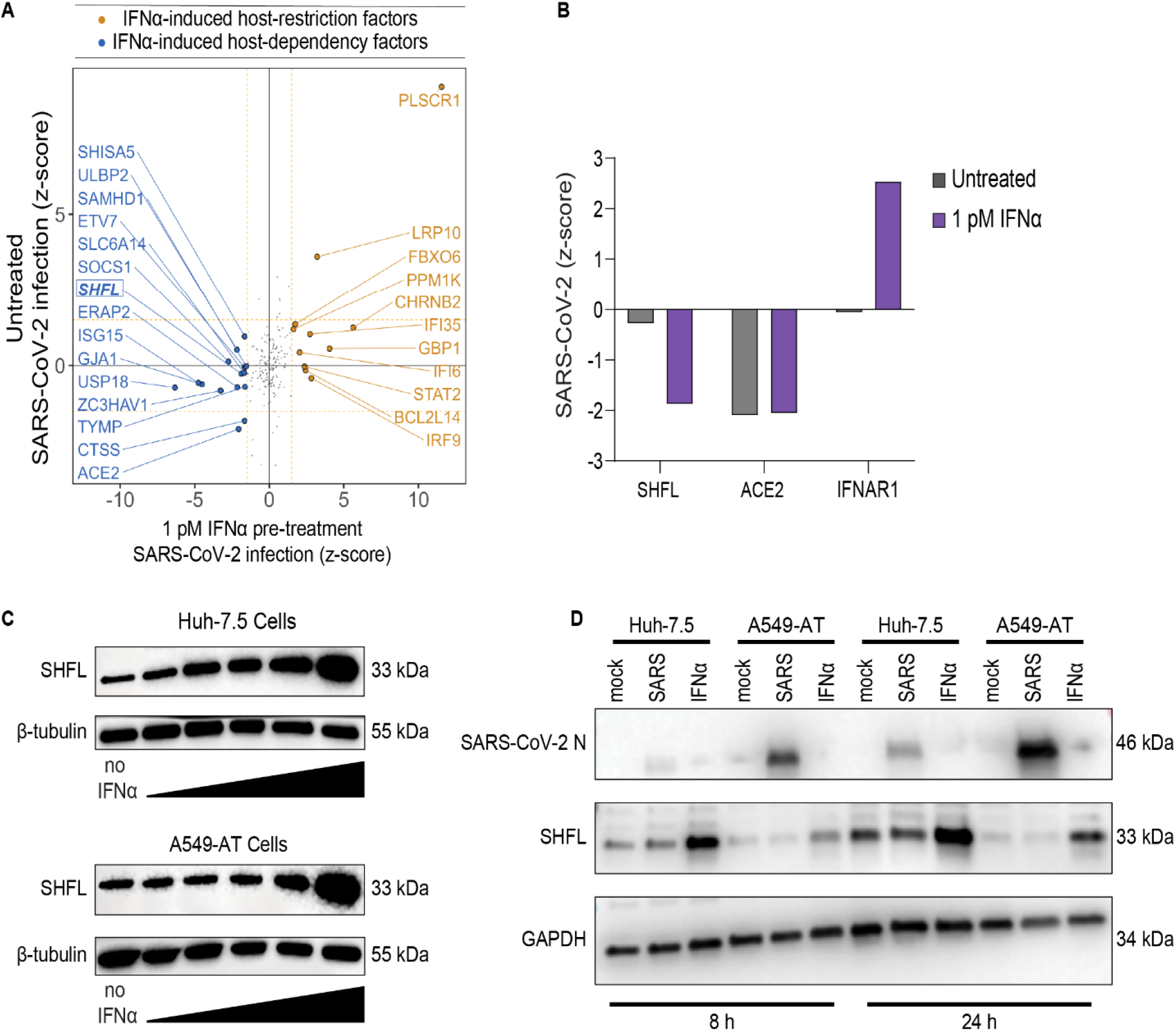
A genome-wide arrayed CRISPR KO screen identifies SHFL as a candidate interferon-dependent host dependency factor for SARS-CoV-2. **(A)** The level of infection (z-score of normalized percent of infected cells) is plotted for 24 h SARS-CoV-2 infection in untreated cells (y-axis) or in cells pre-treated with 1 pM IFNα (x-axis). The plotted genes are a subset identified as ISGs based on mRNA-seq of IFNα-treated cells. Dashed lines mark the z-score cut-off used to call hits (greater than 1.5 or less than −1.5); SHFL is highlighted in bold and outlined. (B) Changes in the SARS-CoV-2 z-score, normalized to NT and total cell count, are plotted for *SHFL* and two gene controls, *ACE2* and *IFNAR1*. Data are represented as a single z-score value that is a composite of all replicates for each gene shown. Data were z-scale normalized with assay plates, then per row and column for spatial effects. Finally, data were Z-scaled across all genes for a final z-score per gene. (C) SHFL inducibility by IFNα in Huh-7.5 and A549-ACE2-TMPRSS2 (A549-AT) cells was measured by Western blotting for the human SHFL protein. SHFL was induced by treating cells with IFNα at increasing concentrations, starting at 0.01 pM and increasing 10-fold to a top dose of 100 pM. Cells were harvested 24 h after treatment. β-tubulin served as a loading control. (D) The ability of SARS-CoV-2 infection alone to induce SHFL expression in Huh-7.5 and A549-AT cells was assessed at 8 and 24 h by Western blotting for human SHFL and the SARS-CoV-2 nucleocapsid protein. Cells were left untreated, infected with SARS-CoV-2, or treated with 100 pM IFNα. Cells were infected at an MOI of 0.01. GAPDH served as a loading control.

Given the novelty of this phenotype and its IFNα-dependence, we characterized baseline SHFL expression across two different SARS-CoV-2 susceptible cell lines in response to exogenous IFNα treatment and SARS-CoV-2 infection. Western blot (WB) analysis of Huh-7.5 and A549-AT cells treated with increasing concentrations of IFNα showed that both cell types express SHFL at baseline, and that SHFL protein induction is dose-dependent (Fig. 1C). SARS-CoV-2 infection alone did not increase SHFL protein expression at 8 h or 24 h post-infection compared with mock-infected cells (Fig. 1D).

Taken together, the screen identifies SHFL as a candidate proviral factor for SARS-CoV-2. While SHFL was IFNα-induced in both Huh-7.5 and A549-AT cells, SARS-CoV-2 alone did not increase expression of SHFL during infection. The remainder of this work focuses on defining the role of SHFL as a host-dependency factor during coronavirus infection.

### Validation of SHFL as a host-dependency factor for alpha- and beta-coronaviruses

To validate SHFL as a potential host-dependency factor for SARS-CoV-2, we generated SHFL KO single-cell clones in Huh-7.5 cells using an mRNA-Cas9 approach. In this method, the Cas9 gene is encoded in an mRNA along with a GFP reporter separated by a P2A site and co-transfected with either SHFL or NT sgRNAs. This allows for transient expression of the Cas9 machinery along with the ability to enrich successfully transfected cells via flow cytometry (Fig. 2A). Cells are sorted via FACS for GFP expression, and only cells in the top 5% of GFP expression levels are retained for single-cell plating and expansion. After enrichment and expansion of candidate SHFL-KO and NT single-cell clones, loss of SHFL protein expression was confirmed by WB analysis. SHFL protein was undetectable in Huh-7.5-SHFL KO single-cell clones under both IFNα pre-treated and untreated conditions, while SHFL expression in Huh-7.5-NT cells remained at baseline and was IFNα-inducible (Fig. 2B). The three single-cell clones for each group shown in panel B were used for all subsequent studies of SHFL-KO cells. After validation of SHFL depletion, SHFL-KO and NT single-cell clones were seeded in both individual and pooled formats in a 1:1:1 ratio of each clone type (to mitigate differences from clonal variation) followed by IFNα pre-treatment and SARS-CoV-2 or HCoV-229E infection. The IFNα concentration used for each virus was selected by completing an IFNα titration for each virus in parental Huh-7.5 cells. The selected concentrations were close to the inhibitory concentration (IC_50_) value for each virus. We selected HCoV-229E for testing alongside SARS-CoV-2 to gain insight into whether SHFL would be a host-dependency factor for coronaviruses outside of the betacoronavirus genus and outside of the highly pathogenic human coronaviruses. HCoV-229E is a seasonal coronavirus belonging to the alphacoronavirus genus that can naturally infect Huh-7.5 cells, making it an ideal candidate in our SHFL KO studies. SHFL KO in Huh-7.5 cells significantly reduced the percentage of SARS-CoV-2-infected cells under untreated and IFNα pre-treated conditions. SHFL KO also reduced infection by the human coronavirus HCoV-229E, but only significantly in the untreated group (Fig. 2C-2E and Fig. S1). Clone-to-clone variability was observed as expected but became negligible through pooling. We then used a focus-forming assay (FFA) to measure the number of infectious virions produced by SARS-CoV-2 or HCoV-229E from SHFL-KO vs NT cells under untreated and IFNα pre-treated conditions. Fewer infectious virions were released by SHFL-KO cells than by NT cells in every group, indicating that the phenotype does not require IFNα pre-treatment (Fig. 2F). However, IFNα pre-treatment increased the magnitude of the effect on released virus. SARS-CoV-2 virion release decreased by 62% in the untreated group but decreased by 89% with IFNα pre-treatment. The same phenotype was observed with HCoV-229E infection with 41% and 74% reduction for untreated and IFNα pre-treated cells, respectively. Overall, these data indicate that the SHFL host-dependency phenotype is not limited to SARS-CoV-2 but extends to a human coronavirus of a second genus, HCoV-229E, and that IFN pre-treatment is not required yet enhances the baseline effect.

**Figure 2.**
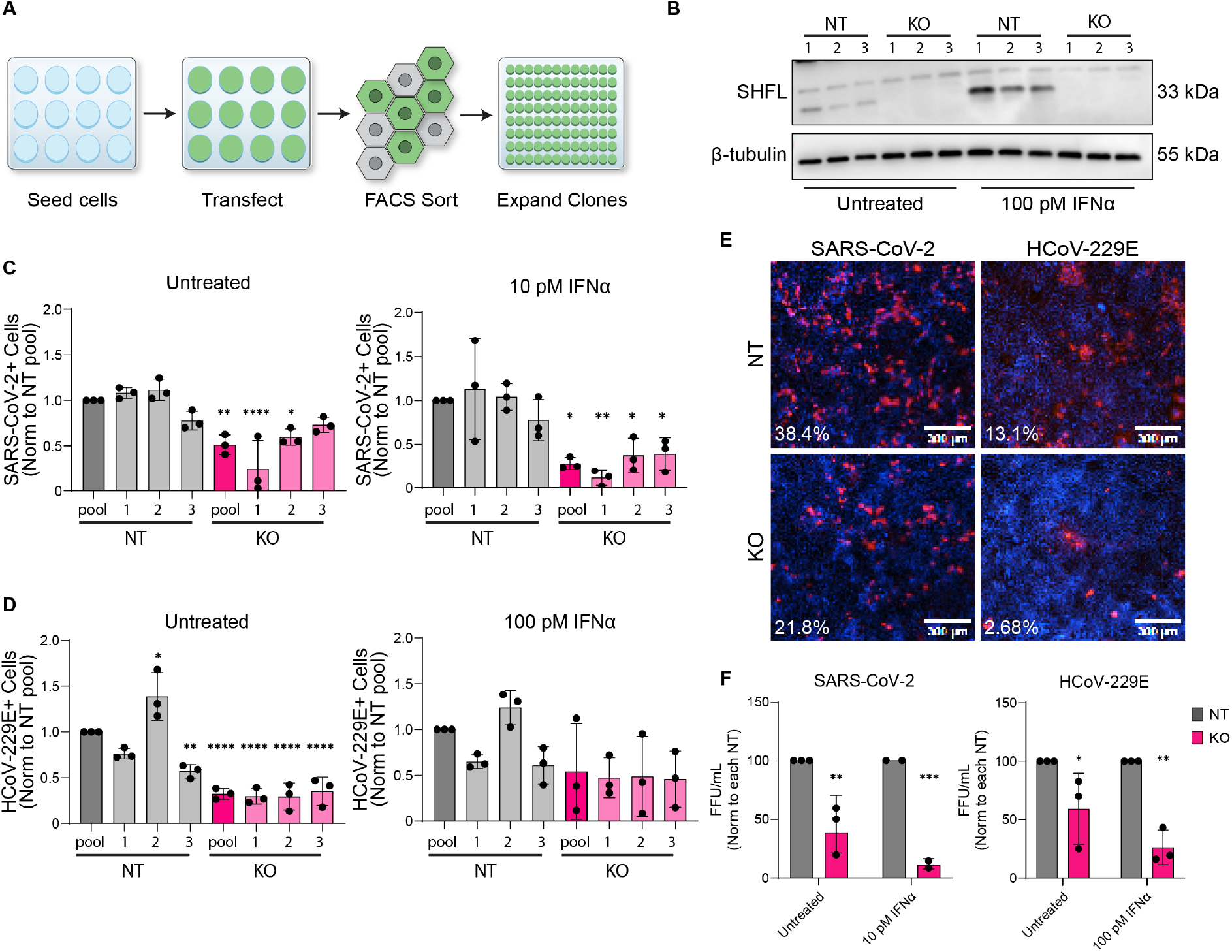
Stable SHFL KO in Huh-7.5 validates host-dependency phenotype of SHFL for SARS-CoV-2 and HCoV-229E. **(A)** Schematic representation of CRISPR-Cas9 KO workflow. Cells were seeded, co-transfected with mRNA-Cas9 and sgRNAs, then FACS sorted for single-cell clone generation. **(B)** SHFL KO efficiency in three Huh-7.5-NT clonal lines and three Huh-7.5-SHFL KO clonal lines was measured by Western blotting for the human SHFL protein. SHFL was induced by treating cells with 100 pM IFNα. Cells were harvested 24 h after treatment. β-tubulin served as a loading control. **(C)** The percentage of total cells infected by SARS-CoV-2 is plotted for Huh-7.5-SHFL KO and Huh-7.5-NT samples in untreated or 10 pM IFNα pre-treated conditions; normalized to the NT pool for each treatment. Cells were seeded either as individual single-cell clones or as a pool of the three clones of each type (1:1:1 ratio). Cells were infected with an MOI of 0.01. Bars show the mean of n = 3 independent experiments, error bars show SD, and each symbol is the average of 4 technical replicates within a single experiment. * P < 0.05, ** P < 0.01, ****, P < 0.0001 (Ordinary one-way ANOVA with Dunnett’s multiple comparisons test). **(D)** The percentage of total cells infected by HCoV-229E is plotted for Huh-7.5-SHFL KO and Huh-7.5-NT samples in untreated or 100 pM IFNα pre-treatment conditions; normalized to the NT pool for each treatment. Cells were seeded either as individual single-cell clones or as a pool of the three clones of each type (1:1:1 ratio). Cells were infected with an MOI of 0.001. Bars show the mean of n = 3 independent experiments, error bars show SD, and each symbol is the average of four technical replicates within a single experiment. ** P < 0.01, ****, P < 0.0001 (Ordinary one-way ANOVA with Dunnett’s multiple comparisons test). **(E)** Representative immunofluorescent images showing the percentage of infected cells for conditions described in Fig. 2C and 2D. Images selected show pooled clonal samples for both SHFL-KO and NT. Cells were stained for either SARS-CoV-2 Nucleocapsid or HCoV-229E Spike (Red) and counterstained with Hoechst nuclear dye (blue). Selected images represent untreated cells only. Scale bar = 300 μm. **(F)** The number of infectious virions / mL in supernatant collected from Huh-7.5-SHFL KO or Huh-7.5-NT cells infected with either SARS-CoV-2 or HCoV-229E was measured in untreated and 10 pM (SARS-CoV-2) or 100 pM (HCoV-229E) IFNα pre-treated conditions. Cells were infected with an MOI of 0.01 for both viruses as measured in Vero-CCL81 cells. Infectious virions were measured using a FFA on parental Huh-7.5 cells. Data are normalized to the Huh-7.5-NT cells within each treatment group. Bars show the mean of n = 3 independent experiments; error bars show SD, and each symbol is the average of four technical replicates within a single experiment. * P < 0.05, ** P < 0.01. *** P < 0.001, ****, P < 0.0001 (Ordinary two-way ANOVA with Sidak’s multiple comparisons test).

### SHFL loss alters basal and IFNα-treated host transcriptional profiles

To investigate whether SHFL influences host cellular processes that may contribute to coronavirus susceptibility, we performed mRNA sequencing of uninfected Huh-7.5-SHFL KO and NT control cells under untreated conditions or following 8 h or 24 h of IFNα treatment. Three independently derived clones per genotype were pooled to minimize clonal variation. Principal component analysis (PCA) revealed separation of samples by IFNα treatment along the first principal component (PC1, 42.1% of total variance), while the second principal component (PC2, 17.6%) distinguished SHFL-KO and NT cells across treatment conditions (Fig. 3A). These findings indicate that SHFL loss is associated with a distinct host transcriptional profile in both untreated and IFNα-treated cells.

**Figure 3.**
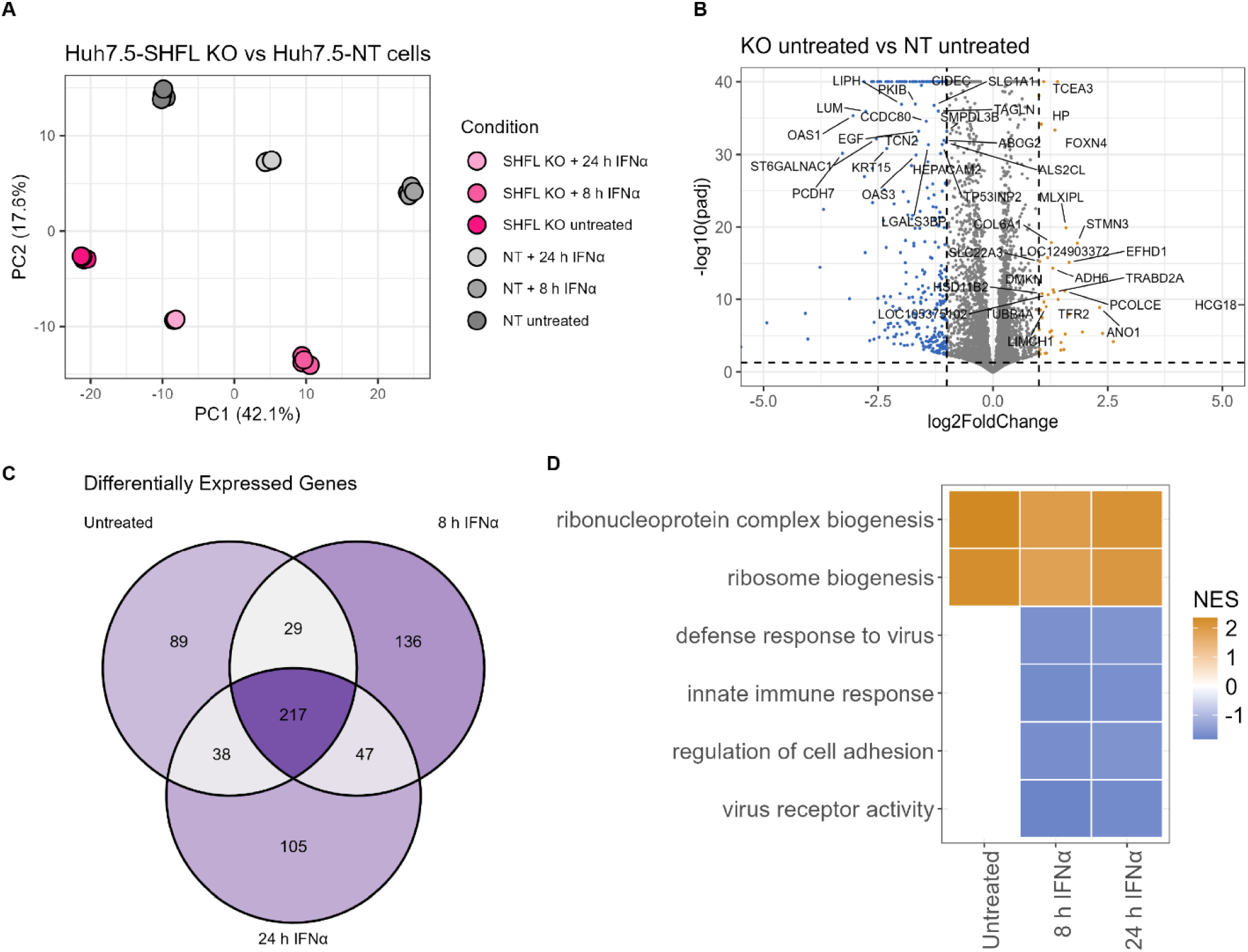
SHFL loss alters basal and IFNα-treated host transcriptional profiles. **(A)** Principal component analysis (PCA) of mRNA sequencing data from Huh-7.5-SHFL KO and Huh-7.5-NT cells under untreated conditions or following treatment with 100 pM IFNα for 8 h or 24 h. Each treatment group contains four replicate samples, and each point represents an individual replicate. **(B)** Volcano plot showing differentially expressed genes (DEGs) in untreated Huh-7.5-SHFL KO cells relative to Huh-7.5-NT controls. Genes were classified as differentially expressed using a Benjamini-Hochberg-adjusted P value < 0.05 and an absolute log2 fold change ≥ 1. Negative log2 fold-change values indicate reduced expression in SHFL-KO cells relative to NT controls, while positive values indicate increased expression. **(C)** Venn diagram showing the overlap of DEGs identified in SHFL-KO versus NT comparisons under untreated, 8 h IFNα-treated, and 24 h IFNα-treated conditions. A total of 217 DEGs were shared across all three comparisons, of which 216 (99.5%) exhibited consistent directionality across conditions. **(D)** Gene set enrichment analysis (GSEA) of Gene Ontology (GO) terms from Biological Process, Molecular Function, and Cellular Component categories, using genes ranked by shrunken log2 fold change for SHFL-KO relative to NT cells within each treatment condition. Six representative pathways were selected from significantly enriched GO terms (Benjamini-Hochberg-adjusted P < 0.05) based on their biological relevance to SHFL function, innate immune responses, and host cellular processes. Tile color represents the normalized enrichment score (NES), with positive values indicating enrichment toward SHFL-KO cells and negative values indicating enrichment toward NT controls. Blank tiles indicate pathways that did not meet the adjusted significance threshold for the corresponding comparison.

To characterize these transcriptional differences, we performed differential gene expression analysis comparing SHFL-KO and NT cells within each treatment condition. Notably, untreated SHFL-KO cells exhibited significant differential expression relative to NT controls (Fig. 3B and Fig. S2), demonstrating that genotype-associated transcriptional differences occur in the absence of viral infection or exogenous IFNα stimulation. Comparison of differentially expressed genes (DEGs) across treatment conditions identified 217 genes shared among all three KO vs NT comparisons, and 216 of these genes changed in the same direction across conditions (Fig. 3C), indicating that SHFL loss is associated with a stable transcriptional program that persists in both untreated and IFNα-treated cells. To characterize the biological processes associated with these transcriptional differences, we performed Gene Ontology (GO) gene set enrichment analysis (GSEA) and selected representative pathways with significant enrichment (BH-adjusted P < 0.05) based on their biological relevance to SHFL function, innate immune responses, and host cellular processes (Fig. 3D). Together, these findings demonstrate that SHFL loss is associated with alterations in host transcriptional programs under basal and IFNα-treated conditions.

### Overexpression studies with SHFL-WT and a SHFL frameshifting domain mutant show that the SHFL proviral phenotype is partially dependent on the frameshifting domain

To test whether this novel host-dependent phenotype for SHFL is cell-type specific and whether it is linked to the same domain responsible for SHFL’s known antiviral mechanism of inhibiting −1 PRF, we used a gain-of-function approach in A549-AT cells, a human alveolar adenocarcinoma cell line, which overexpress the SARS-CoV-2 receptor ACE2 and co-receptor TMPRSS2. Unlike Huh-7.5 cells, these cells are more physiologically relevant for the study of respiratory viruses and have more intact innate immune responses. Plasmids overexpressing the canonical SHFL isoform (SHFL-WT), a naturally occurring isoform in which the known −1PRF domain from amino acids 164-199 is deleted (SHFL-Δ; RefSeq NP_001295206.1), or an Empty control were transduced into A549-AT cells (Fig. 4A)^39,44^. Overexpression of SHFL-WT and SHFL-Δ was validated by WB for human SHFL protein with GAPDH as a loading control. Notably, the size difference between SHFL-WT and SHFL-Δ matches the domain deletion (Fig. 4B). Cell viability was assessed by measuring metabolic activity using the resazurin-based PrestoBlue assay. The PrestoBlue assay showed no significant differences between the three cell lines (Fig. S3). To test whether SHFL-Δ cells retain proviral activity compared with SHFL-WT and Empty controls, cells were seeded in 96-well plates and left untreated or pretreated with 10 pM IFNα. At 24 h post-treatment, cells were infected with SARS-CoV-2 for 16 h. SHFL-Δ attenuates the SHFL-WT phenotype under both untreated and treated conditions. Under IFNα pre-treatment SHFL-WT reached significance against Empty controls while SHFL-Δ did not; neither reached significance in the untreated condition (Fig. 4C).

**Figure 4.**
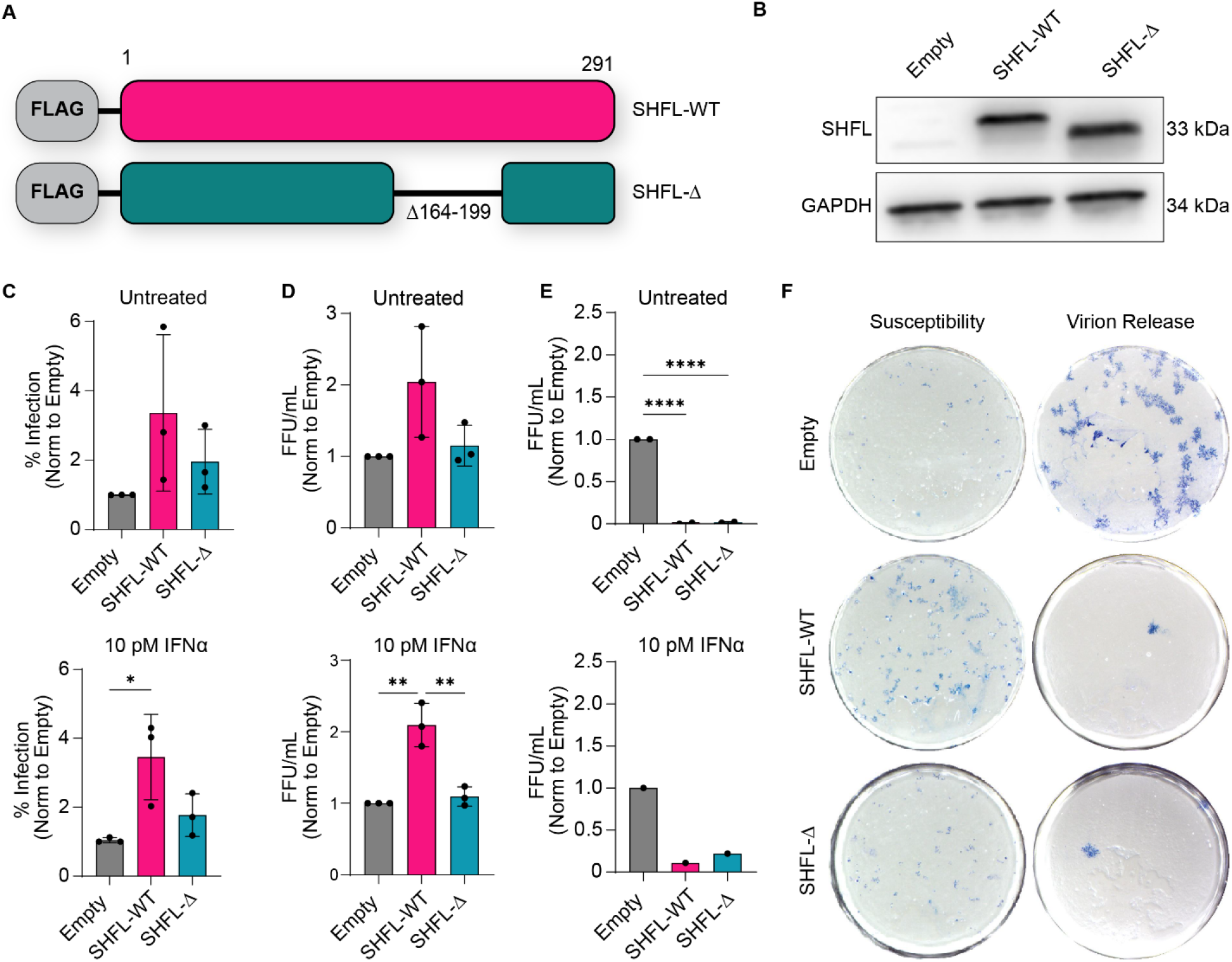
Overexpression of SHFL-WT and SHFL frameshift domain mutant in A549-AT cells reveal SHFL increases SARS-CoV-2 susceptibility but decreases virion production. **(A)** Schematic representation of SHFL constructs used for overexpression experiments. SHFL-WT and SHFL-Δ were cloned into the pWPI vector. Each SHFL gene contains an N-terminal 3xFLAG tag. SHFL-Δ contains a deletion from amino acids 164-199 corresponding to the known antiviral frameshifting domain. The Empty pWPI vector was used as a control. (B) SHFL overexpression was measured in A549-AT-Empty, A549-AT-SHFL-WT, and A549-AT-SHFL-Δ cells by Western blotting for human SHFL protein. GAPDH served as a loading control. (C) The percentage of total cells infected by SARS-CoV-2 as measured by immunofluorescent staining for Nucleocapsid protein is plotted for A549-AT-Empty, SHFL-WT, and SHFL-Δ overexpressing cells in untreated or 10 pM IFNα pre-treated conditions. SARS-CoV-2 infection was measured by immunofluorescent staining for SARS-CoV-2 Nucleocapsid. Each bar is normalized to the % of cells infected in the Empty control condition. Cells were infected with an MOI of 0.01 for 16 h. Bars show the mean of n = 3 independent experiments, error bars show SD, and each symbol is the average of four technical replicates within a single experiment. * P < 0.05 (Ordinary one-way ANOVA with Tukey’s multiple comparisons test). (D) Susceptibility of SHFL-overexpressing cells to SARS-CoV-2 infection was measured with a direct FFA. Empty, SHFL-WT, and SHFL-Δ cells were seeded and either left untreated or pre-treated with 10 pM IFNα for 24 h. Viral foci were quantified using a SARS-CoV-2 Nucleocapsid antibody. Bars show the mean of n = 3 independent experiments, error bars show SD, and each symbol is the average of four technical replicates within each experiment. ** P < 0.01 (Ordinary one-way ANOVA with Tukey’s multiple comparisons test). (E) SARS-CoV-2 particle production from infected SHFL-overexpressing cells was measured by FFA using infected supernatants. Empty, SHFL-WT, and SHFL-Δ cells were seeded and pre-treated with either 0 pM or 10 pM IFNα for 24 h. Cells were infected with an MOI of 0.01, and the infected supernatants were collected at 72 hours post-infection (hpi). Bars show the mean of n = 2 for untreated and n = 1 for 10 pM IFNα treated, error bars show SD, and each symbol is the average of four technical replicates within each experiment. **** P < 0.0001 (Ordinary one-way ANOVA with Tukey’s multiple comparisons test). (F) Representative images of SARS-CoV-2 foci under the FFA conditions described in Fig. 4D and Fig. 4E. Cells were stained for SARS-CoV-2 N protein followed by incubation with HRP-tagged secondary antibody and detection with SeraCare TrueBlue Peroxidase substrate.

Our first questions to define the proviral mechanism were: how does SHFL affect early susceptibility to virus and how does SHFL affect the release of infectious virions. The susceptibility of SHFL-WT and SHFL-Δ cells to SARS-CoV-2 infection under IFNα-untreated and pre-treated conditions was tested using a FFA assay. SHFL-WT showed a two-fold increase in SARS-CoV-2 susceptibility compared to SHFL-Δ and Empty controls with or without IFNα pre-treatment (Fig. 4D and 4F). To test the effects of SHFL on virion release, we performed a FFA on supernatants collected from SARS-CoV-2-infected SHFL-WT or SHFL-Δ cells. Supernatants from infected SHFL-WT and SHFL-Δ cells showed a 99% decrease in infectious virus compared to Empty controls (Fig. 4E and 4F). This reduction therefore did not require the −1 PRF domain, in contrast to the hypothesis above. Overall, these data show that the ability of SHFL to enhance SARS-CoV-2 infection is not cell-type specific, and that SHFL-WT both increases susceptibility to SARS-CoV-2 and reduces the number of infectious virions released.

### SHFL decreases sensitivity of SARS-CoV-2 to IFNα independent of the frameshifting domain

To test whether SHFL-overexpressing cells are preferentially infected relative to Empty control cells, and whether any preference depends on IFN signaling, we used a mixed-population approach. To isolate the effect from IFN signaling we used a combination of IFNα and ruxolitinib treatments. Ruxolitinib is a JAK/STAT inhibitor and was used to block JAK/STAT signaling from IFNα treatment in our cells. We expect that any effects of SHFL-WT on SARS-CoV-2 infection that require JAK/STAT signaling would be eliminated after IFNα treatment. To conduct the experiment, SHFL-WT and Empty cells were mixed in a 50:50 ratio during seeding. These cells were treated with or without 10 μM ruxolitinib, a saturating dose of the inhibitor, for 1 h, followed by treatment with or without 10 pM IFNα. The cells were infected with SARS-CoV-2 24 h after IFNα treatment (Fig. 5A). To determine whether SHFL-overexpressing cells were preferentially infected by SARS-CoV-2, we used IF staining to quantify the number of infected cells (SARS-CoV-2 N) and the number of SHFL-overexpressing cells (FLAG). Analysis of the percentage of FLAG+ cells showed that the true baseline composition of each well was around 60% SHFL-WT cells, rising to around 70% after IFNα treatment (Fig. S4). Of the total infected cells, we calculated the proportion that were SHFL-overexpressing and that were Empty control cells. A value equal to the number of FLAG+ cells in the group would indicate no preference for SHFL-overexpressing cells in SARS-CoV-2 infection. The average percentage of infected cells that overexpress SHFL was 76% in untreated cells, indicating that SHFL-overexpressing cells are more likely to be infected by SARS-CoV-2 than Empty controls. Interestingly, when this mixed population was pre-treated with IFNα, the proportion of infected cells that were SHFL-overexpressing increased (84%) (Fig. 5B and 5C). Combined with the SHFL KO data in Figure 1, which showed a larger phenotype in IFNα-pretreated cells, we hypothesized that SHFL may negatively regulate the IFN response to enhance SARS-CoV-2 infection, in addition to increasing SARS-CoV-2 susceptibility. To test this hypothesis, SHFL-WT, SHFL-Δ cells, or Empty control cells were pre-treated with a serial dilution of IFNα for 24 h before SARS-CoV-2 infection. The percentage of infected cells was measured by IF staining for SARS-CoV-2 N protein, and these values were used to generate IC_50_ curves for IFNα in each cell line (Fig. 5D). Both A549-AT-SHFL-WT and A549-AT-SHFL-Δ cells showed an increase in the IC_50_ of IFNα compared to Empty control (Table 1). We observed a similar effect in the Huh-7.5-SHFL KO cells. The IC_50_ in SHFL KO cells decreased compared to NT but the data was challenging to replicate without significant standard deviation rates due to the low overall infection levels in Huh-7.5-SHFL KO cells. To confirm that this effect is specific to IFN signaling, we used ruxolitinib to block the effects of IFNα. To do this, SHFL-WT, SHFL-Δ, and Empty cells were pre-treated with increasing concentrations of ruxolitinib for 1 h, followed by treatment with 10 pM IFNα. The cells were again infected with SARS-CoV-2 24 h after IFNα treatment, and infection was measured by IF staining (Fig. 5E). The ability of ruxolitinib to rescue SARS-CoV-2 infection levels in the SHFL-WT and SHFL-Δ cells was blunted compared to Empty controls. Infection in Empty cells increased an average of 3.3-fold compared to baseline IFNα treatment but only increased 1.7- and 2.3-fold, respectively, in SHFL-WT and SHFL-Δ cells (Table 2). These findings suggest that the reduced sensitivity to IFNα observed in SHFL-WT and SHFL-Δ cells may involve negative regulation of canonical JAK/STAT signaling. Overall, this supports our hypothesis that SHFL also enhances SARS-CoV-2 infection by reducing the sensitivity of cells to IFNα and suggests that this phenotype does not require the previously characterized frameshifting domain.

**Figure 5.**
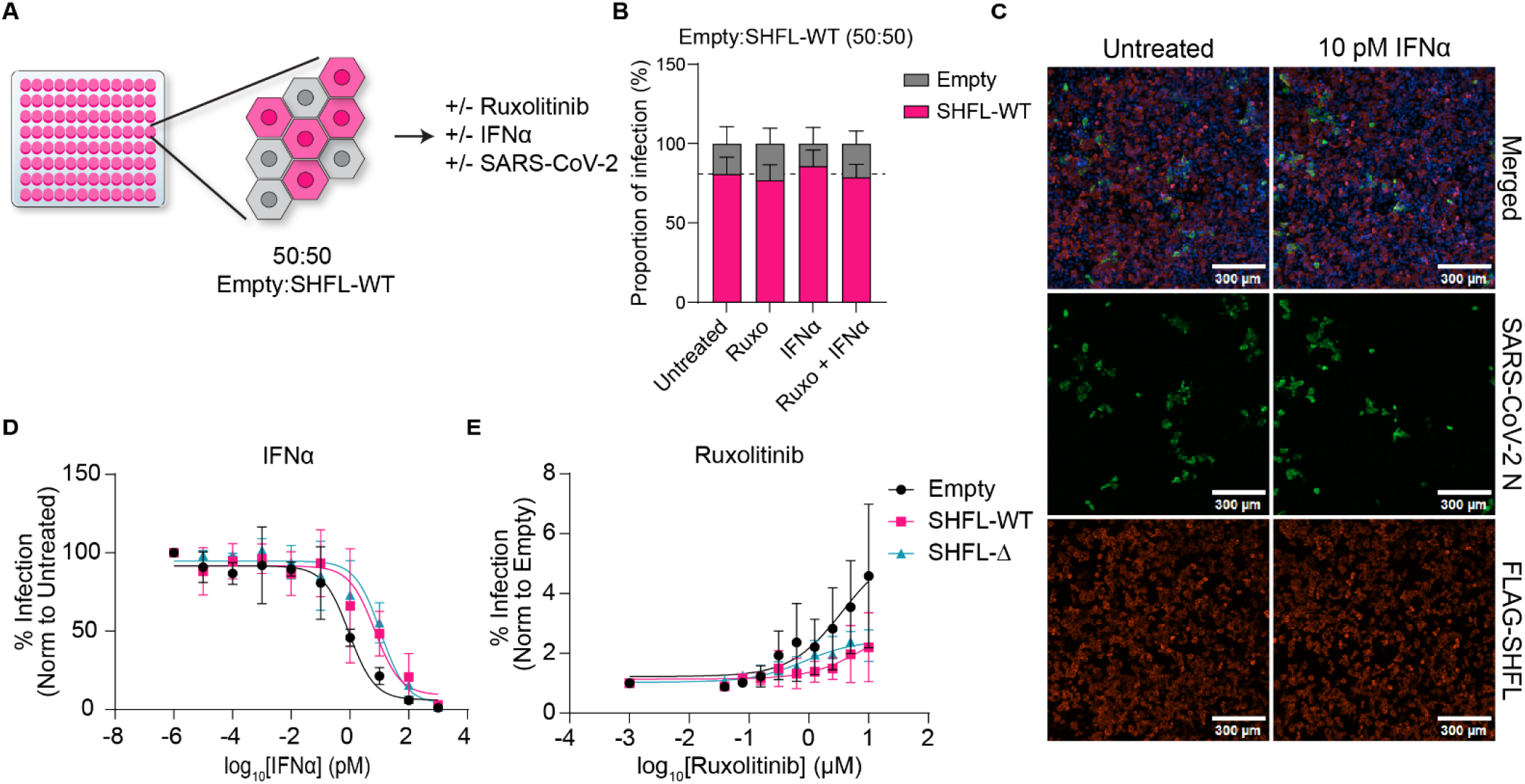
SHFL decreases SARS-CoV-2 sensitivity to IFNα independent of the frameshift domain. **(A)** Schematic representation of mixing experiments used to determine probability of infection for A549-AT-SHFL-WT cells compared to Empty controls. A549-AT-Empty and A549-AT-SHFL-WT cells were mixed in a 50:50 ratio at the time of seeding followed by ruxolitinib and/or IFNα pre-treatment and SARS-CoV-2 infection. **(B)** The percentage of the number of infected cells that were SHFL-overexpressing vs Empty control is plotted for the cell mixing experiment outlined in Fig. 5A. Cells were either untreated, treated with 10 μM of ruxolitinib alone, 10 pM of IFNα alone, or 10 μM of ruxolitinib followed by 10 pM IFNα treatment. SARS-CoV-2 infection and SHFL overexpression were measured by immunofluorescent staining for SARS-CoV-2 N protein and FLAG, respectively. Cells were infected with an MOI of 0.01. Bars show the mean of n = 3 independent experiments, and error bars show SD. Each individual experiment is represented by the average of four technical replicates. ** P < 0.01 (Repeated Measures two-way ANOVA with Sidak’s multiple comparisons test). **(C)** Representative immunofluorescent images showing the percentage of infected cells that were SHFL-overexpressing as described in Fig. 5B. Cells were stained for SARS-CoV-2 N (Green), FLAG-tag (Red), and counterstained with Hoechst nuclear dye (Blue). Selected images represent cells left untreated or pretreated with 10 pM IFNα. Scale bar = 300 μm. **(D)** IC_50_ curves for the inhibitory capacity of IFNα pre-treatment against SARS-CoV-2 infection in A549-AT-Empty, A549-AT-SHFL-WT, and A549-AT-SHFL-Δ are shown. Cells were pre-treated with IFNα at decreasing concentrations, starting at 1000 pM and decreasing 10-fold to 0.00001 pM. After 24 h, the cells were infected with SARS-CoV-2 at an MOI of 0.01. Each point represents an average of n = 3 independent experiments, error bars = SD. **(E)** EC_50_ curves for the effectiveness of ruxolitinib at blocking IFNα activity against SARS-CoV-2 are shown for A549-AT-Empty, A549-AT-SHFL-WT, and A549-AT-SHFL-Δ cells. Cells were pre-treated with decreasing concentrations of ruxolitinib, starting at 10 μM and two-fold to 0.15 μM. After 1 h cells were treated with 10 pM IFNα. 24 h after IFNα treatment, cells were infected with SARS-CoV-2 at 0.01 MOI. Each point represents an average of n = 4 independent experiments, error bars = SD.

**Table 1.** Susceptibility of SARS-CoV-2 to IFNα in A549-AT SHFL-overexpressing cells. Values represent the average and SDs from n = 3 independent experiments. Bolded values in parentheses indicate fold change in IC_50_ compared to A549-AT-Empty cells. Each experiment was composed of three technical replicates per condition.

| **IC<sub>50</sub> ± SD (fold change from Empty)** |  |
| **Cell Line** | IFNα (pM) |
|---|---|
| **A549-AT-Empty** | 1.134 ± 0.1021 **(1)** |
| **A549-AT-SHFL-WT** | 9.738 ± 6.472 **(8.6)** |
| **A549-AT-SHFL-Δ** | 11.91 ± 8.985 **(10.5)** |

**Table 2.** Susceptibility of SARS-CoV-2 infection in IFNα pre-treated A549-AT SHFL overexpressing cells to ruxolitinib treatment. Shown are the effective concentration (EC_50_) values and max effect size for ruxolitinib on SARS-CoV-2 in cells pre-treated with 10 pM IFNα. Max effect size indicates the fold change in the percent of SARS-CoV-2 infected cells in the highest ruxolitinib dose (10 μM) compared to untreated cells within each cell line. Values represent the average and SDs from n = 3 independent experiments. Bolded values in parentheses indicate fold change compared to A549-AT-Empty cells. Each experiment was composed of three technical replicates per condition.

| **Max Effect Size and EC<sub>50</sub> ± SD (fold change from Empty)** |  |  |
| **Cell Line** | Ruxolitinib (µM) | Max Effect Size |
|---|---|---|
| **A549-AT-Empty** | 0.5342 ± 0.2262 **(1)** | 3.363 ± 1.476 **(1)** |
| **A549-AT-SHFL-WT** | 0.7137 ± 0.4091 **(1.4)** | 1.697 ± 0.7273 **(0.5)** |
| **A549-AT-SHFL-Δ** | 0.5079 ± 0.3504 **(0.95)** | 2.377 ± 0.2581 **(0.7)** |

## Discussion

Here, we show that the interferon-stimulated gene SHFL, long characterized as an antiviral restriction factor, also acts as a host-dependency factor for human coronaviruses. SHFL knockout in Huh-7.5 cells reduced infection by SARS-CoV-2 and by the seasonal alphacoronavirus HCoV-229E; conversely, overexpression of SHFL in A549-AT cells increased susceptibility to SARS-CoV-2. In a mixed-population assay, SARS-CoV-2 preferentially infected SHFL-overexpressing cells, and that preference increased after IFNα pre-treatment. SHFL-overexpressing cells were also less sensitive to IFNα, an effect that may involve negative regulation of canonical JAK/STAT signaling and did not appear to require the previously characterized −1 PRF domain. Together, these results place SHFL among the growing group of ISGs with both antiviral and proviral activity.

SHFL is best known as an inhibitor of −1 PRF during SARS-CoV-2 and HIV-1 replication. SHFL is a *bona fide* RNA-binding protein that binds both RNA and translational machinery to restrict −1 PRF during translation of viral and cellular genes^45^. Whether −1 PRF inhibition itself is sufficient or required for that antiviral activity is, however, already contested: Jager et al. reported that inhibition of −1PRF is dispensable against HIV-1 and that SHFL also restricts murine leukemia virus (MLV), which does not use −1 PRF^46^. Supporting this finding, Wang et al. reported that phosphorylation of residues T250 and T253, outside the defined −1 PRF domain (AA164-199), is essential for antiviral activity against HIV-1^41^. He et al. placed −1 PRF inhibition under zinc-dependent control of SHFL’s N-terminal autoinhibition^35^. Against that background, a proviral activity that does not map to the −1 PRF domain is plausible, and that is what we observe: deleting AA164-199 leaves the effect of SHFL on IFNα sensitivity intact.

In our study, we used two cell line models to examine SHFL’s effect on coronavirus infections. Our choice of cell lines followed this rationale. We selected Huh-7.5 cells for the initial screening and characterization because of their specific defects in innate immune signaling. These have defective RIG-I functionality and produce little IFNα in response to SARS-CoV-2 infection, while remaining responsive to exogenous IFNα (Fig. 1D)^34,47^. That combination provides a clean background for ISG studies: infection itself cannot drive induction of SHFL or of other ISGs, so we can use these cells to measure the phenotype of ISG knockouts at baseline endogenous levels with and without IFN signaling and downstream ISG expression. A549-AT cells were selected for our overexpression and mechanistic studies due to their physiological relevance for SARS-CoV-2 infection and allowed us to test the SHFL proviral phenotype across multiple cell lines.

Although SHFL has primarily been characterized as an antiviral effector, emerging evidence suggests that it also contributes to cellular processes outside the context of viral infection. Kelly and Dinman recently implicated SHFL in ribosome stress surveillance, proposing that its constitutive expression contributes to the recognition of aberrantly frameshifted ribosomes^36^. Consistent with a broader cellular role for SHFL, our transcriptomic analysis revealed differences between SHFL-KO and NT cells even in the absence of viral infection or exogenous IFNα stimulation. The strong directional concordance of shared DEGs across untreated and IFNα-treated conditions suggests that SHFL loss establishes a reproducible host transcriptional state rather than producing only treatment-specific transcriptional effects. Notably, ribonucleoprotein complex biogenesis was consistently enriched in SHFL-KO cells across untreated and IFNα-treated conditions, suggesting that SHFL loss is associated with changes in host RNA-associated processes. However, whether these transcriptional differences arise directly from altered ribosome surveillance or reflect secondary cellular responses remains unknown. Our analysis also identified differences in immune-associated transcriptional programs, including reduced expression of several antiviral-associated genes in SHFL-KO cells following IFNα treatment. Interestingly, these findings do not readily explain the reduced coronavirus susceptibility observed in SHFL-KO cells, as increased expression of canonical antiviral effectors would instead be expected to favor viral restriction. Together, these observations suggest that the transcriptional changes associated with SHFL loss do not provide a straightforward explanation for its proviral activity. Further investigation of transcriptional responses during infection and the functional consequences of these pathways will be necessary to establish their contribution to SHFL-mediated coronavirus susceptibility.

Here we report, in both gain-of-function and loss-of-function studies, that SHFL can promote infection of SARS-CoV-2 in two distinct human cell lines and of the seasonal coronavirus HCoV-229E in Huh-7.5 cells. In our initial screen, SHFL KO in Huh-7.5 cells showed an IFN-dependent host-dependency phenotype. However, follow-up experiments in stable Huh-7.5-SHFL KO cells showed that this IFN signaling is not required for the baseline host-dependency phenotype, but that it can enhance the magnitude of the response. This highlights the importance of screening validation. Screening is advantageous for hit identification but is not always accurate mechanistically due to confounding variables. In our screen, we had no metric to assess overall SHFL KO efficiency, but our stable cell lines showed no detectable SHFL protein by WB validation. This background difference could explain the differences between our screening and validation results. The lack of significance for the decrease in HCoV-229E infection in SHFL KO cells pre-treated with IFNα (Fig. 2D) could reflect a lower overall infection rate or confounding by viral spread. Overall, these data indicate that the SHFL host-dependency phenotype is not limited to SARS-CoV-2 but extends to multiple human coronaviruses from different genera of the *Coronaviridae* family and that this phenotype shows IFN-independence. The low baseline level of SHFL is sufficient to promote viral infection, but the IFN response further enhances it by inducing additional SHFL expression. This phenotype has been seen before but not characterized further. In a study of SHFL’s antiviral activity against flavivirus infection, Hanners et al. reported in an overexpression screen for multiple viruses and virus families in Huh-7.5 cells stably overexpressing SHFL that SHFL increased infection with both SARS-CoV-2 and the seasonal coronavirus HCoV-OC43, providing support for our results^43^.

To further characterize how SHFL promotes coronavirus infection, we used a SHFL mutant lacking the defined −1 PRF domain, SHFL-Δ, to examine whether this domain plays a role in both antiviral and proviral functions. Upon infection initiation, SHFL-WT increases cell susceptibility to SARS-CoV-2, and this phenotype is attenuated in SHFL-Δ cells, indicating its involvement, but loss of this domain alone is not sufficient to restore SARS-CoV-2 infection to baseline. In addition, SARS-CoV-2 infection is further enhanced in SHFL-WT overexpressing cells pre-treated with IFN, and SHFL can also decrease the sensitivity of cells to IFNα, resulting in reduced control of the viral infection. The effect of ruxolitinib, a JAK/STAT inhibitor, was also reduced upon IFNα treatment in SHFL-WT and SHFL-Δ cells. Together, these findings suggest that SHFL modulates cellular sensitivity to IFNα through a mechanism involving JAK/STAT signaling that is independent of the previously characterized −1 PRF domain. However, whether SHFL directly alters downstream IFN-stimulated transcription remains unknown. Future studies using an interferon-stimulated response element (ISRE) reporter assay could determine whether SHFL expression affects IFN-induced transcriptional activity and further clarify its role in regulating the IFN response.

Our two principal observations suggest opposing effects of SHFL on SARS-CoV-2 infection. SHFL-WT roughly doubled the susceptibility of A549-AT cells to SARS-CoV-2, yet supernatants from those same cells yielded 99% less infectious virus than Empty controls. These findings suggest that SHFL promotes infection establishment while restricting subsequent production of infectious progeny. Unlike overall infection which showed a proviral phenotype across both Huh-7.5 loss-of-function and A549-AT gain-of-function experiments, the effect on virion production differed between the two cell lines. This could be due to differences in virus spread or other factors between the two cell models. Additional experiments distinguishing intracellular viral replication, assembly, and virion release will be needed to identify the stage responsible for reduced infectious-virus production. Interestingly, deletion of the previously characterized −1 PRF domain attenuated the observed increase in viral susceptibility but did not eliminate the reduction in infectious-virus output. This suggests that the latter phenotype may involve an antiviral mechanism independent of the deleted domain, consistent with previous evidence that SHFL-mediated restriction of HIV-1 does not necessarily require −1 PRF inhibition^46^.

As there is a growing body of evidence that ISGs have multiple mechanistic functions, both antiviral and proviral, we propose that this novel mechanism of SHFL is just one of multiple mechanisms involved during coronavirus infection. Understanding more about the complex nature of virus-host interactions with innate immune molecules brings us one step closer to understanding how we can better leverage innate host defenses against future emerging viruses.

## Methods

### Plasmids, oligos, and primers

Plasmids encoding wild-type human SHFL and the frameshift-domain mutant human SHFL with N-terminal FLAG epitope tags were a gift from V. Kinast.

sgRNAs for SHFL CRISPR KO, including non-targeting controls, are from Horizon Discovery (SG-020478-01-002, SG-020478-02-002, U-007501-01-20, U-002005-20).

### Cell Lines

Huh-7.5 (human hepatocellular carcinoma; *H. sapiens*; sex: male^34^; Dr. Charles M. Rice), A549-ACE2-TMPRSS2 (human lung carcinoma; *H. sapiens*; sex: male; generously provided by the lab of Dr. Charles Rice)^31^, Lenti-X 293T (*H. sapiens*; sex: female; Takara, cat. #632180), Vero-CCL81 (*C. aethiops* kidney epithelial cells; sex: female; generously provided by the lab of Dr. Raymond Schinazi), and Vero-E6 cells (*C. sabaeus* kidney epithelial cells; sex: female; generously provided by the lab of Dr. Vineet Menachery) were cultured in Dulbecco’s Modified Eagle Medium supplemented with 10% fetal bovine serum (FBS) and 1% nonessential amino acids (NEAA) at 37°C with 5% CO_2_. Cells are routinely screened for mycoplasma and have tested negative.

### Virus Stocks

SARS-CoV-2: The genome-wide arrayed CRISPR KO screen was infected using virus stocks as described previously^26^. For the remaining experiments, icSARS-CoV-2-nLuc was acquired from BEI resources (NR-54003). To generate icSARS-CoV-2-nLuc stocks, Vero-CCL81 cells were infected at 80% confluence with 0.001 MOI of P0 icSARS-CoV-nLuc. Virus was harvested at 48 h post-infection and clarified by centrifugation at 2000 x g for 10 min at 4°C. Virus was tittered on Vero-CCL81 cells using a focus-forming assay as described below. SARS-CoV-2 infections for the genome-wide arrayed CRISPR KO screen were conducted as previously described^26^. SARS-CoV-2-nLuc infections were performed using infection media (DMEM + 2% FBS + 1% NEAA) with an absorption time of 60 min at the indicated MOI.

HCoV-229E: HCoV-229E was acquired from BEI resources (NR-52726). Huh-7.5 cells were used to propagate and titer the virus as described in the previous paragraph. HCoV-229E infections were performed using infection media (DMEM + 2% FBS + 1% NEAA) with an absorption time of 90 min at the indicated MOI.

### Virus Concentration

SARS-CoV-2 was concentrated via centrifugation using Amicon Ultra Centrifugal Filters with a 100kDa molecular weight cut-off. Virus was concentrated 10X at 1000 x g, 4°C in 15-30 min increments. Virus was not concentrated unless otherwise noted in the manuscript text.

### Unbiased Genome-Wide Arrayed CRISPR KO Screening

The genome-wide arrayed CRISPR KO screen was conducted as previously reported in Le Pen, Paniccia et al. The raw data were adapted for this publication to highlight SHFL specifically but were not otherwise altered^26^.

### Western Blotting

Cell lysates were prepared in RIPA buffer supplemented with 1:100 of Halt Protease and Phosphatase Inhibitor Cocktail and thawed on ice. Lysates were clarified by centrifugation at 16,000 × g for 10 min at 4°C. Protein concentrations were measured using the Pierce BCA kit. Samples were prepared for gel electrophoresis by combining with 3X loading dye and 10X DTT reagents (NEB; B7703S) before denaturing at 75°C for 10 min. Samples were run on 4-20% 10-well or 15-well Mini-Protean TGX protein gels (Biorad; 4561096) at 200 V for 40 min. Proteins were transferred onto nitrocellulose membranes using the Trans-Blot Turbo Transfer System (Biorad; 1704158EDU). Ponceau S red dye was used to ensure complete transfer of proteins to the membrane, followed by 3 washes with TBS/T (1 X TBS + 0.1% Tween-20) and incubation for 1 h at room temperature in blocking buffer (TBS/T + 5% milk). Membranes were incubated with primary antibody diluted in blocking buffer overnight at 4°C (SHFL, abcam, ab122765, 1:500; β-tubulin, Proteintech, HRP-66240, 1:10,000; SARS-CoV-2 Nucleocapsid, GeneTex, GTX135357, 1:2,000; GAPDH, Invitrogen, MA1-16757, 1:1000). Membranes were washed 3 x with TBS/T and then incubated with either HRP-anti-mouse or HRP-anti-rabbit secondary antibodies (Invitrogen; 31430 and 31460). Membranes were developed with the Immobilon Forte Western HRP substrate (Millipore; WBLUF0500) and were imaged using the Azure600. To complete secondary and tertiary rounds of antibody staining, original antibodies were removed by incubating membranes in Restore Western Blot Stripping buffer (Thermofisher; FER21059X4) for 30 min at room temperature, followed by 3 washes with 1 X TBS. Membranes were then re-blocked and probed as described above.

### Immunofluorescent (IF) staining

Cells were fixed with 4% PFA followed by three washes with PBS. Cells were permeabilized by incubating with PBS + 0.1% Triton X-100 at room temperature for 10 min. Cells were washed three times with PBS, then incubated with blocking buffer (PBS + 5% normal goat serum) for 1 h at room temperature. Primary antibodies were diluted in blocking buffer and added to the cells overnight at 4°C (SARS-Cov-2 N, GeneTex, GTX135357, 1:2000; HCoV-229E S RBD, R&D Systems, MAB10938, 1:1000; FLAG, Proteintech, 66008-4-Ig, 1:2000). Cells were then washed three times with PBS + 0.1% Tween-20 solution. Secondary antibodies and Hoechst dye counterstain were diluted in blocking buffer and incubated on the cells for 1 h at room temperature (Hoechst, Thermofisher, 33342, 1:5000; AlexaFluor 488 goat anti-rabbit, ThermoFisher; cat no. A-11034, 1:2000; AlexaFluor 594 goat anti-mouse, Thermofisher, A-11032, 1:2000). Cells were washed three times with PBS and stored at 4°C until imaging.

### Cytation7 immunofluorescent imaging and analysis

The Cytation7 Cell Imaging Multi-Mode Reader and Gen5 software (Agilent) were used to image and quantify all immunofluorescent staining experiments. Plates were imaged at 4x unless otherwise noted. Analysis and cell counting were performed by first quantifying all Hoechst+ nuclei to determine the cell number. 594 and 488 fluorophores were used to identify which sub-populations of the Hoechst+ cells were positive for viral antigens and host genes. Positivity thresholding for infected or SHFL-overexpressing cells was determined by identifying the mean fluorescent intensity for a given fluorophore in mock infected or empty vector wells and setting the minimum threshold at 3 standard deviations above the mean. The cellular mask to determine positivity was extended 5 μm outside the boundaries of the nucleus to include cytoplasmic staining. Mean fluorescent intensities and positive cell populations were recorded for all fluorescent channels used.

### Focus-Forming Assay + ELIspot staining

96-well plates were coated with collagen from calf skin for 1 h. Cells were seeded at 10k cells per well in DMEM + 10% FBS + 1% NEAA. Cells were infected with either SARS-CoV-2 or HCoV-229E in serial dilution (absorption times listed above) followed by a methylcellulose overlay (DMEM + 1% methylcellulose + 2% FBS). Plates were fixed with 4% PFA after 48 h. Plates were permeabilized and blocked as described above. Primary antibody (SARS-CoV-2 N or HCoV-229E S RBD) was diluted in blocking buffer and applied to cells overnight at 4°C. Cells were washed and incubated with HRP-conjugated secondary antibody diluted in blocking buffer. Plates were washed, dried, and incubated with KPL TrueBlue peroxidase substrate (SeraCare) for 1 h at room temperature. Plates were dried and imaged with the Immunospot CTL machine.

### mRNA-Cas9 CRISPR KO transfection

Huh-7.5 cells were co-transfected with 1.75 ng/μL of mRNA-Cas9 (Horizon Discovery) and either 25nM NT crRNA:tracrRNA or 25nM SHFL sgRNA (1:1 ratio of sgRNA1 and sgRNA2; Horizon Discovery) using 0.4% DharmaFECT Duo transfection reagent (Horizon Discovery). Cells were enriched for single-cell clone generation via FACS sorting at 24 h post-transfection.

### Lentivirus Production

Lenti-X 293T cells were seeded in T175 flasks. At 80% confluency, the cells were co-transfected with 5.5 μg of psPAX2 and 2 μg of pMD2.G, and 7.5 μg of the delivery plasmid using 45 μL of FuGENE transfection reagent (Promega; cat. no. E5911). Plasmids were diluted in 683 μL of Opti-MEM reduced serum medium. Media was changed at 16 h post-transfection (hpt) and lentiviruses were harvested at 48 hpt. Lentiviruses were clarified by centrifugation at 500 x g, 4°C for 10 min to remove cell debris.

### Lentivirus Transduction

Cells were seeded for lentivirus transduction in 6-well plates at 100k cells per well or 96-well plates at 10k cells per well. One hour prior to transduction, cells were changed to low-serum media (DMEM + 2% FBS + 1% NEAA). Lentiviruses were diluted in low-serum media to the desired concentration along with 4 μg/mL of polybrene to enhance transduction efficiency. Cells were spinoculated for 1 h at 1000 x g, 37°C. The lentivirus inoculum was replaced with 10% DMEM (DMEM + 10% FBS + 1% NEAA) at 16 h post-transduction.

### Generation of SHFL-overexpressing cells

pWPI-SHFL-WT, pWPI-SHFL-Δ, or pWPI-Empty plasmids were delivered to A549-AT cells via lentiviral transduction. Lentiviruses were initially titrated on A549-AT cells to determine a dilution for each lentivirus resulting in 10-30% transduction efficiency to prevent multiple transduction events per cell. Cells were transduced with the selected dilution of each lentivirus in 6-well plates. Positively transduced cells were selected with Blasticidin selection media (DMEM + 10% FBS + 1% NEAA + 5 μg/mL Blasticidin) at 2 days post-transduction before expansion.

### Fluorescent Activated Cell-Sorting (FACS)

FACS sorting was performed with support from the Emory Pediatrics/Winship Flow Cytometry Core. Cells were dissociated from their plates via trypsinization. Cells were resuspended in 10% DMEM, filtered through a 0.22 μm strainer, and centrifuged (1200 x g, 4°C, 5 min). Media was removed, cells were resuspended with PBS to wash and centrifuged again (1200 x g, 4°C, 5 min). Each sample was then resuspended in 1 mL unsupplemented DMEM. Cells were filtered again into 5 mL FACS tubes (BD 352063). Cells were sorted on the BD FACSAria II and gating was determined using the BD FACSDiva 9.4 software. Cells were gated for live cells, followed by singlets, then GFP+ expression. The top 5% of GFP+ cells were selected for single-cell plating into 96-well plates containing 100 μL of conditioned media per well (DMEM + 10% FBS + 1% NEAA collected from parental cells, clarified via centrifugation (1200 x g, 4°C, 5 min) and then filtered with a 0.22 μm filter).

### mRNA-seq

#### Cell Culture and IFNα treatment

Approximately 200k Huh-7.5-NT or Huh-7.5-SHFL KO cells were seeded in each well of a 6-well plate with 2 mL of DMEM + 10% FBS + 1% NEAA media. The next day, cells were either left untreated or treated with 100 pM IFNα. Cells were harvested at either 8 h or 24 h post-treatment in 1 mL TRIzol (Invitrogen; cat no. 15596-018. N = 4 replicates (separate wells) per sample.

#### RNA Extraction

Each 1 mL TRIzol sample was loaded into MaXtract High Density tubes (2 mL, Qiagen, 129056). 200 μL of chloroform was added to each tube. The tubes were vortexed for 20 s for thorough mixing. Samples were rested at room temperature for 2-3 min after vortexing. After resting, the MaXtract tubes were centrifuged at 12,000 x g for 10 min at 4°C. The aqueous layer at the top of the MaXtract tubes (approx. 600 μL) was transferred to RNase-free tubes pre-filled with 600 μL of ethanol. Ethanol and aqueous mixture were vortexed thoroughly. Sample was transferred to columns from the RNeasy Mini Kit (Qiagen; cat no. 74104). From there, the manufacturer’s protocol was followed to complete the RNA extraction including the option on-column DNase digestion after washing with buffer RW1 (Qiagen DNase I; cat no. 79254). RNA was eluted with RNase-free water and stored at −80°C.

#### Sequencing

mRNA sequencing was performed by Novogene. Purified total RNA was subjected to poly(A) enrichment for library preparation. Libraries were sequenced on an Illumina NovaSeq X Plus platform using 150 bp-length reads. Read quality was evaluated with FastQC v0.11.9 prior to, and following, trimming^48^. Adapters were removed and low-quality reads filtered with fastp v1.3.1, retaining only reads with a mean quality of at least Q20 and a length of at least 36 bp^49^. Transcript abundances were then estimated from the trimmed reads with Salmon v1.11.4 in selective-alignment mode against a partial decoy-aware index built from the GENCODE v49 human transcriptome, with the GRCh38 primary assembly included as decoy^50^. Gene-level counts were obtained by summarizing the transcript-level estimates with tximport in R^51^.

#### mRNA-seq Differential Expression and Pathway Analysis

mRNA-seq gene counts were analyzed in R using the DESeq2 package^52^. Genes were filtered to those with ≥ 10 counts. Differential expression was assessed using pairwise contrasts between Huh-7.5-SHFL KO and Huh-7.5-NT cells for each treatment condition and within cell-line comparisons between treatment timepoints. Log_2_FC estimates were shrunk using the adaptive shrinkage estimate implemented in the ashr package to improve stability of effect-size estimates for genes with low counts or high dispersion^53^. Differential expression results were visualized as volcano plots using the ggplot2 package with genes classified according to the adjusted *P* value and log_2_ fold-change thresholds (adjusted *P* < 0.05, and |log_2_ fold-change| ≥ 1).

Gene set enrichment analysis (GSEA) was performed using the ClusterProfiler package with Gene Ontology (GO) annotations from org.Hs.eg.db^54^. Genes were ranked by shrunken log_2_fold-change for each comparison, and GO terms from Biological Process, Molecular Function, and Cellular Component ontologies were analyzed using the gseGO function. Gene sets containing 3–800 genes were included, and statistical significance was corrected for multiple testing using the Benjamini-Hochberg method^54^. The 20 GO terms with the smallest adjusted *P* values across each comparison group were combined, and normalized enrichment scores (NES) were visualized using ggplot2^55^.

#### mRNA-seq principal-component and PC2 loading analysis

For principal-component analysis (PCA), RNA-seq counts were variance-stabilized using the DESeq2 variance-stabilizing transformation. PCA was performed using the prcomp function. Sample scores along PC1 and PC2 were visualized by experimental condition. Gene contributions to PC2 were obtained from the PC2 loading coefficients, and the 25 genes with the largest absolute PC2 loading values were visualized. For pathway analysis of PC2-associated genes, Ensembl-IDs were converted using AnnotationDbi and the org-Hs-eg-db database^56,57^. All mapped genes were ranked according to their signed PC2 loading coefficients and subjected to preranked GSEA using the fgsea package with the Hallmark gene sets from Molecular Signatures Database (MSigDB)^58,59^.

### Cell Mixing Experiment

A549-AT-Empty and A549-AT-SHFL-WT were mixed in a 1:1 ratio before seeding in 96-well plates. Cells were treated with 10 μM ruxolitinib or DMSO control for 1 h followed by ± 10 pM IFNα treatment for a total of four distinct treatment arms. Cells were infected with SARS-CoV-2 at an MOI of 0.01 for 16 h before fixing for immunofluorescent staining and analysis.

### IFNα IC_50_ Curve Generation

Cells were seeded in 96-well plates followed by treatment with serial dilution of IFNα in DMEM + 10% FBS + 1% NEAA. The starting dilution was 1000 pM and decreased 10-fold to 0.00001 pM. After 24 hours, cells were infected with SARS-CoV-2 at an MOI of 0.01.

### Ruxolitinib EC_50_ Curve Generation

Cells were seeded in 96-well plates followed by treatment with serial dilution of ruxolitinib in DMEM + 10% FBS + 1% NEAA + 0.1% DMSO. Starting concentration of ruxolitinib was 10 μM and decreased 2-fold to 0.15 μM.

## Acknowledgements

We thank the members of the Michailidis, Schinazi, and Sarafianos labs for discussion of the data. We thank Dr. Charles M. Rice for the Huh-7.5 and A549-AT cells and Dr. Vineet Menachery for the Vero-E6 cells.

This work was supported by the Emory University Research Committee (URC) Award for the Biological and Health Sciences and the Emory University Molecules and Pathogens to Populations and Pandemics Seed Grant Award (MP3). This research was supported in part by the National Institutes of Health (R01 AI167356-01A1 to S.G.S.). The content is solely the responsibility of the authors and does not necessarily represent the official views of the National Institutes of Health. S.G.S. acknowledges the Nahmias-Schinazi Distinguished Chair in Research.

## Author Contributions

Taylor M Shue: conceptualization, methodology, software, validation, formal analysis, investigation, writing – original draft, writing – review and editing, visualization, funding acquisition; Georgios Mourtzouchos: Validation, investigation, writing – review and editing; Dar-Yin Li: Investigation, writing – review and editing; Yee T. Ong: Investigation, writing – review and editing; Antonis Athanasiadis: Investigation, writing – review and editing. Maria Bousali: Methodology, formal analysis, software, data curation, writing – editing and review; Madeleine K. Sanders: Investigation, writing – review and editing; Sophie Spector: Software, visualization, writing – review and editing; Christopher River-Bonilla: investigation, writing – review and editing; Khadija Ndoye: Investigation, writing – review and editing; Kenneth C Levenson: Resources, writing – review and editing; Jens Cuba: Investigation, resources, writing – review and editing; Jayden Harris: Investigation, writing – review and editing; Kelly Mieck: Methodology, resources, writing – review and editing; Shuiyun Lan: Methodology, resources, writing – review and editing; Volker Kinast: Methodology, resources, writing – review and editing; Julia LeCher: Methodology, investigation, resources, writing – review and editing; Jérémie Le Pen: Conceptualization, methodology, software, resources, writing – review and editing, supervision; Stefan G. Sarafianos: Methodology, resources, writing – review and editing, supervision, funding acquisition; Eleftherios Michailidis: Conceptualization, methodology, validation, resources, writing – original draft, writing – review and editing, supervision, project administration, funding acquisition.

## Data Availability

mRNA-sequencing data are deposited in the NCBI Sequence Read Archive (SRA) under BioProject accession PRJNA1544138. All other relevant data are within the paper and the associated figures.

---

**Supplemental Figure 1.**
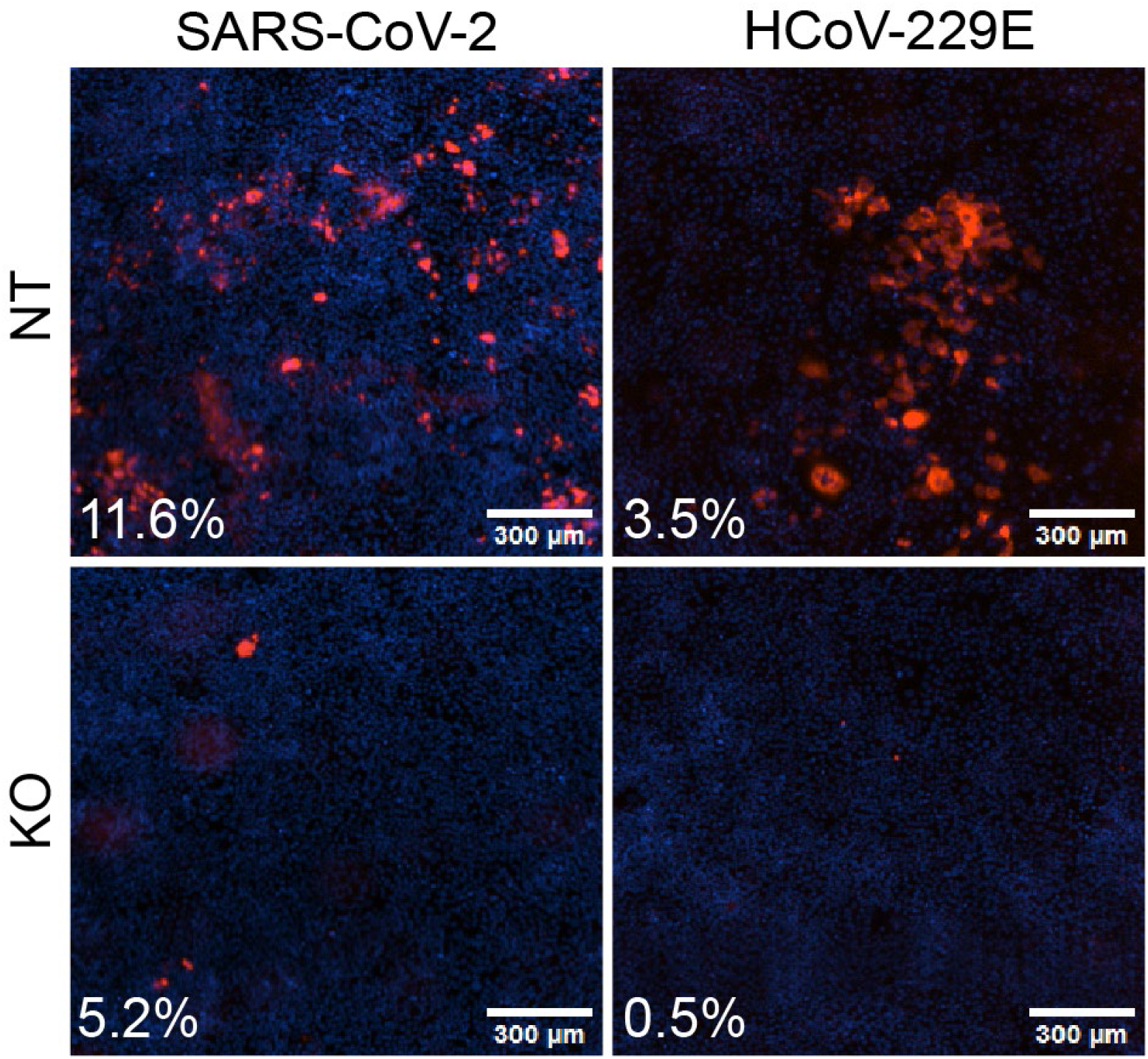
SARS-CoV-2 and HCoV-229E infection in Huh-7.5-SHFL KO cells with IFNα pre-treatment. Representative immunofluorescent images showing the percentage of infected cells for conditions described in Fig. 2C and 2D. Cells were stained for either SARS-CoV-2 N protein or HCoV-229E Spike (red) and counterstained with Hoechst nuclear dye (blue). Selected images represent cells pre-treated with 10 pM (SARS-CoV-2) or 100 pM (HCoV-229E) IFNα only. Scale bar = 300 μm.

**Supplemental Figure 2.**
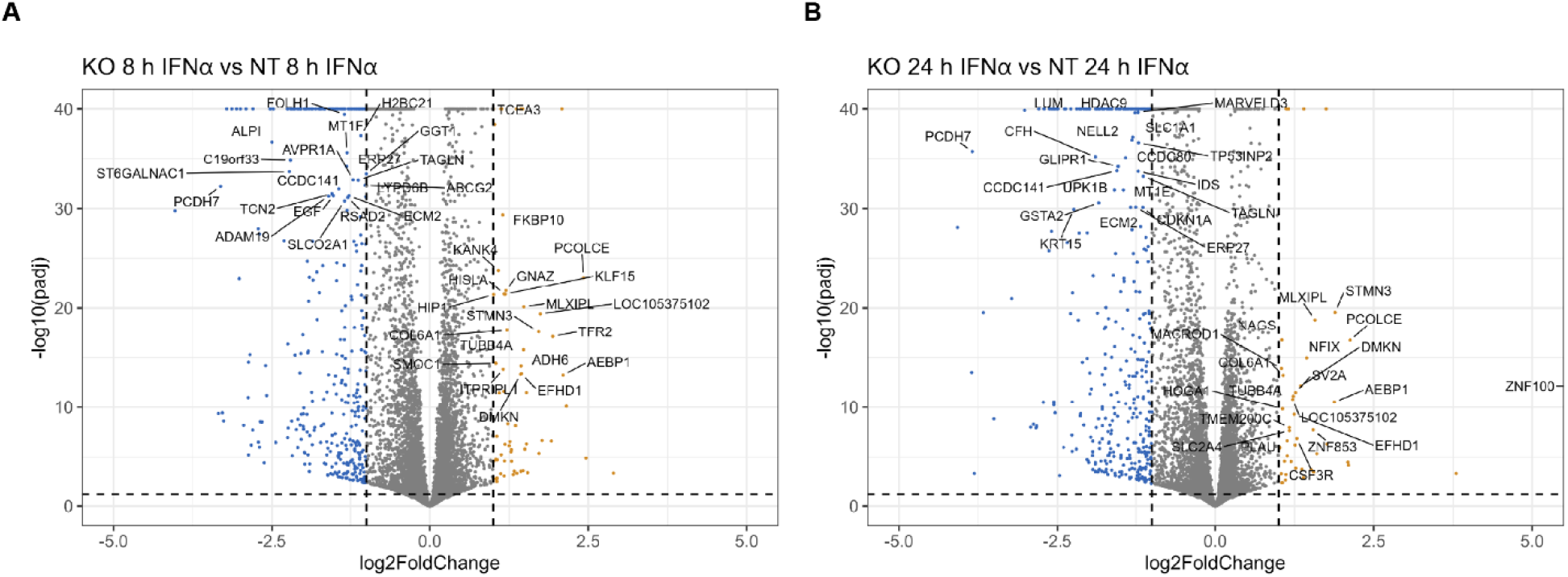
SHFL KO alters the gene expression profiles compared to NT controls. **(A)** Differentially expressed genes in Huh-7.5-SHFL KO and Huh-7.5-NT cells under 100 pM IFNα treatment for 8 h are shown as a volcano plot comparing log_2_ fold change (FC) in gene expression vs log_10_ adjusted p-value. Log_2_FC >= 1 and adjusted p-value < 0.05 were considered significant. Differential gene expression analysis was completed using the DESeq2 package in R. −log_10_(adjusted p-values) were capped at 40 and the top 20 genes are displayed. **(B)** Differentially expressed genes in Huh-7.5-SHFL KO and Huh-7.5-NT cells under 100 pM IFNα treatment for 24 h are shown as a volcano plot comparing log_2_ fold change (FC) in gene expression vs log_10_ adjusted p-value. Log_2_FC >= 1 and adjusted p-value < 0.05 were considered significant. Differential gene expression analysis was completed using the DESeq2 package in R. −log_10_(adjusted p-values) were capped at 40 and the top 20 genes are displayed. For all plots, gold represents genes significantly upregulated in SHFL-KO cells and blue represents genes significantly downregulated in SHFL-KO cells. Grey indicates genes that are not significantly changed in SHFL-KO vs NT cells.

**Supplemental Figure 3.**
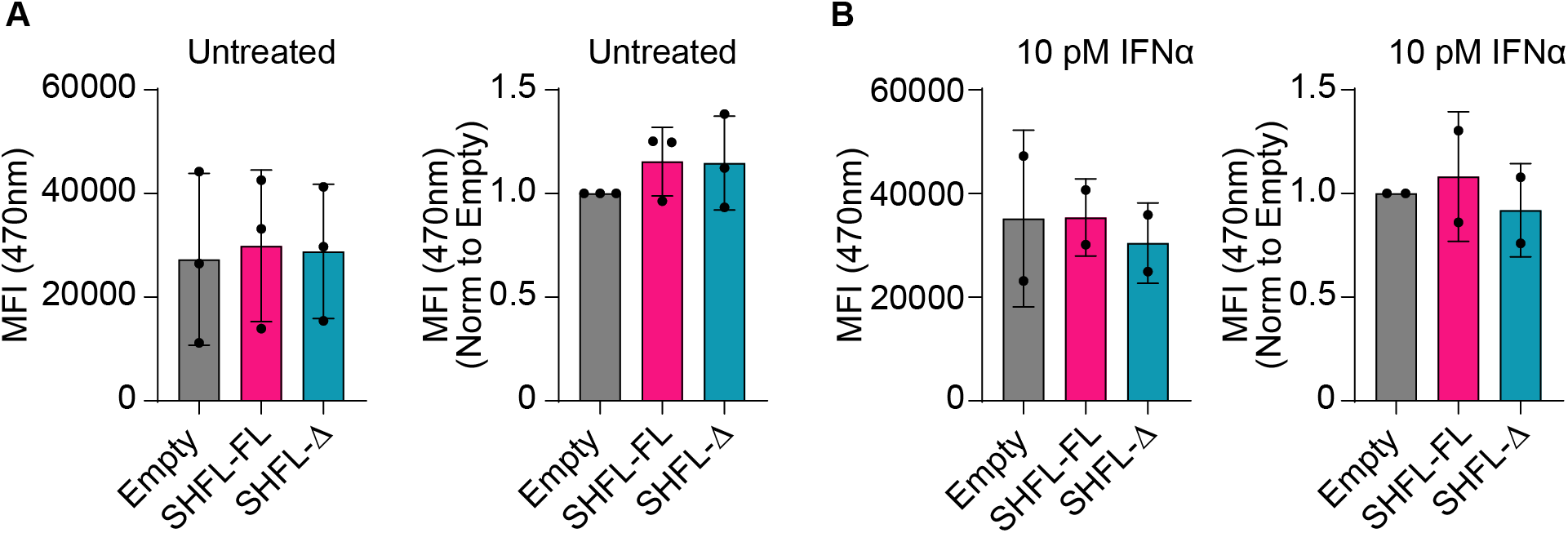
SHFL overexpression in A549-AT cells does not alter cell viability. **(A)** The mean fluorescent intensity (MFI) at 470nm resulting from the PrestoBlue resazurin-based assay is shown for A549-AT-Empty, −SHFL-WT, and SHFL-Δ overexpressing cells in untreated conditions. Bars show the mean of n = 3 independent experiments, error bars show SD, and each symbol is the average of five technical replicates within a single experiment. Data are not significant. (Ordinary one-way ANOVA with Tukey’s multiple comparisons test). **(B)** The mean fluorescent intensity (MFI) at 470nm resulting from the PrestoBlue resazurin-based assay is shown for A549-AT-Empty, −SHFL-WT, and SHFL-Δ overexpressing cells under 10 pM IFNα treatment conditions. Bars show the mean of n = 2 independent experiments, error bars show SD, and each symbol is the average of five technical replicates within a single experiment. Data are not significant. (Ordinary one-way ANOVA with Tukey’s multiple comparisons test).

**Supplemental Figure 4.**
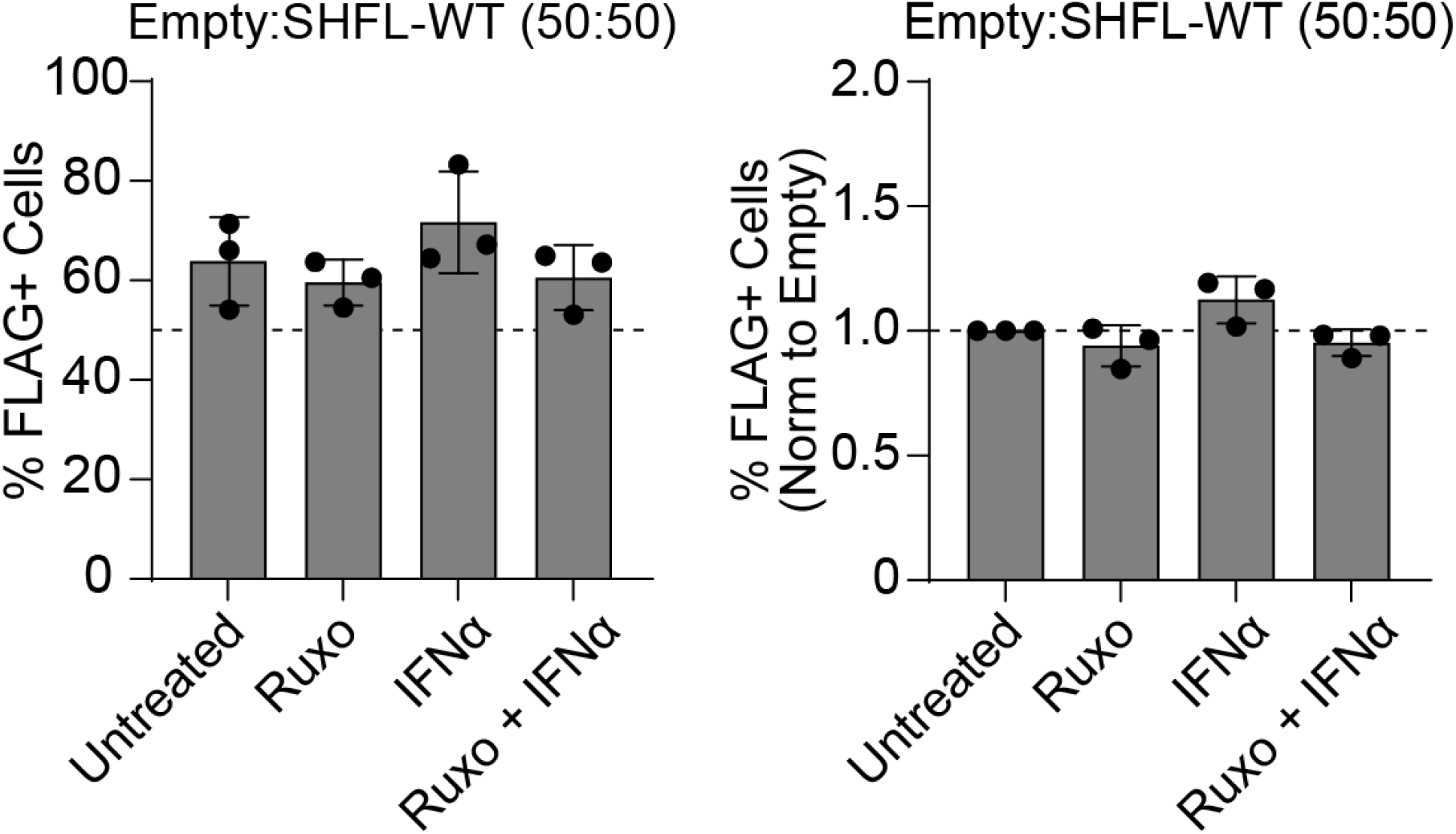
SHFL-overexpression rate in A549-AT-Empty and A549-AT-SHFL-WT mixing experiment. The percentage of SHFL-overexpressing cells in each condition, as measured by immunofluorescent staining, is shown for the experiment described in Fig. 5A and 5B. Cells were stained for the FLAG-tagged SHFL construct. Bars represent the mean of n = 3 independent experiments; error bars show SD. Each symbol represents the average of four technical replicates per experiment. Data are shown as both the raw percentage of FLAG+ cells and the percentage of FLAG+ cells normalized to the untreated group. Data are not significant. (Repeated measures one-way ANOVA with Sidak’s multiple comparisons test).

